# *Mycobacterium tuberculosis* disease associates with higher HIV-1-specific antibody responses

**DOI:** 10.1101/2022.12.02.518812

**Authors:** Bukola Adeoye, Lydia Nakiyingi, Yvetane Moreau, Ethel Nankya, Alex J. Olson, Mo Zhang, Karen R. Jacobson, Amita Gupta, Yukari C. Manabe, Mina C Hosseinipour, Johnstone Kumwenda, Manish Sagar, AIDS Clinical Trials Group A5274 (REMEMBER) Study Team

## Abstract

*Mycobacterium tuberculosis* (Mtb) can enhance immune responses against unrelated pathogens. Although Mtb is the most common co-infection in people living with HIV (PWH), there has been no examination of its impact on HIV-1 immune responses. Plasma neutralization and antibody dependent cellular cytotoxicity (ADCC) was compared among PWH and Mtb disease (PWH/Active Mtb) and PWH/No Mtb both prior to and after antiretroviral treatment (ART) and completion of Mtb therapy. We assessed HIV-1 sequences, total antibody quantities and isotypes, and plasma cytokine levels to ascertain mechanisms that affect humoral responses. HIV-1 neutralizing antibodies (nAbs) were broader and more potent in PWH/Active Mtb as compared to PWH/No Mtb, and nAbs increased among PWH who developed Mtb after ART initiation. ADCC was also higher in the PWH who had Mtb disease after starting ART. PWH/Active Mtb as compared to PWH/No Mtb had unique HIV-1 envelope sequence motifs associated with neutralization resistance further implying differences in humoral selection. The Mtb-linked antibody augmentation associated with elevated plasma cytokine levels important for B cells and antibody production, namely interleukin-6, a proliferation-inducing ligand (APRIL), and B-cell activating factor (BAFF). Increased plasma virus levels, greater HIV-1 envelope diversity, higher levels of all antibodies, and cross-reactive responses did not explain the enhanced HIV-1 humoral responses in those with Mtb. Mtb disease enhances HIV-1 humoral responses likely by perturbing pathways important for antibody production in lymphoid tissue that has both pathogens. These findings have implications for using antibody-based therapies and inducing optimal HIV-1 antibody responses.

**Author Summary:** Mycobacterium tuberculosis (Mtb) is the most common infection among people with HIV (PWH) in the world. Mtb infection can enhance immune responses against unrelated pathogens. Previous studies have not examined the impact of Mtb disease on HIV antibodies in PWH. This information has importance for future strategies aimed at enhancing HIV antibody responses in naïve individuals or PWH. We show that HIV neutralizing antibodies and antibody-dependent cellular cytotoxicity are broader and more potent in PWH in the presence as compared to the absence of Mtb disease. PWH and Mtb disease as compared to those without Mtb also harbor unique HIV envelope sequences, which further indicates that there is differential antibody selection pressure. The Mtb linked HIV antibody enhancement associated with specific mediators important for B cell and antibody development. Importantly, the Mtb mediated HIV antibody augmentation was not due to cross-reactivity, a generalized increase in all antibodies, or a higher level, more diverse, or longer duration of antigen exposure. We speculate that more potent HIV antibodies arise in lymphatic tissue that harbors both Mtb and HIV. Our findings have implications for both future uses of HIV antibodies as prophylaxis or treatment and strategies aimed inducing better HIV antibody responses.

## Introduction

Heterologous immunity describes immune responses induced against a pathogen or an antigen by an unrelated stimulus (1). For instance, a previous viral infection can elicit beneficial or pathologic cross-reactive responses after exposure to another related but different virus (2, 3). Similarly, viral infections can also induce autoimmunity because of cross-reactivity between a virus epitope and a self-antigen (4). Beyond cross-reactivity, heterologous immunity can occur through a phenomenon termed ‘trained immunity’ when one exposure, classically mycobacterial infections, induce epigenetic changes that enhance innate immune responses against a subsequent unrelated infection (5, 6). Mycobacteria, specifically *Mycobacterium bovis* bacillus Calmette-Guerin (BCG), induced trained immunity provides protection against secondary influenza, yellow fever virus, and malaria infections in human clinical trials (5,7,8).

Other mycobacteria, such as *Mycobacterium tuberculosis* (Mtb), may stimulate immune responses against unrelated pathogens by other means besides trained immunity, such as bystander cell activation. Mtb infection imparts diffuse immune activation, which subsequently influences pathways associated with antibody production, potency, and functionality (2). For example, heat-inactivated Mtb in oil, termed complete Freund’s adjuvant (CFA), constitutes one of the most powerful activators of the humoral immune response (9, 10). CFA promotes antigen uptake and processing in antigen-presenting cells. This triggers a cytokine storm that impacts cellular and antibody responses (10, 11). CFA’s use is banned in humans and highly restricted in animals, and thus there is limited ability to examine its impact in a prospective manner. Interestingly, people with human immunodeficiency virus type 1 (HIV-1) with prior Mtb exposure have lower plasma virus levels as compared to those with no Mtb infection, possibly implying an immune interaction (12). While immunization with Mtb and BCG antigens has been shown to promote higher antibody levels against HIV-1 and simian immunodeficiency virus (SIV) proteins in animal models (13, 14), there is limited human evidence for the impact of Mtb on HIV-1 humoral responses.

Two billion people in the world are infected with Mtb (15), and Mtb is the most common co-infection in people with HIV-1 (PWH) (16). While the majority of Mtb infected individuals have latent “inactive” infection, untreated HIV-1 accelerates the development of Mtb disease (17). Furthermore, antiretroviral treatment (ART) initiation can restore anti-Mtb immune responses and “unmask” previously hidden Mtb disease (18). While there are extensive studies of the impact of HIV-1 and ART on Mtb, surprisingly, the influence of Mtb on HIV-1 immune responses remains poorly characterized. We and others have previously demonstrated that PWH and Mtb disease have a unique inflammatory profile that can potentially impact the humoral response (19, 20). For instance, increased interleukin (IL)-6 observed in PWH and Mtb disease can drive the secretion of IL-21 in naïve and memory T cells to promote antibody production (21). Mycobacteria and HIV-1 can also co-exist in the same anatomic region, such as lymph nodes, and even in the same cell, such as macrophages (16,22,23), and this co-localization may also impact the subsequent immune response. Based on CFA’s known actions, unique Mtb-induced inflammation, and Mtb-HIV-1 co-localization, we hypothesized that Mtb disease may augment humoral immune responses against HIV-1.

Here, we show that PWH with as compared to those without active Mtb disease have broader and more potent HIV-1 humoral responses. The augmented HIV-1 antibody response is associated with specific plasma mediators known to be important for antibody production. Importantly, the enhanced antibody response is not due to the non-specific induction of all antibodies or cross-reactivity. These findings suggest that active Mtb disease induces heterologous humoral immunity against HIV-1 likely by perturbing antibody production pathways in tissues that harbor both HIV-1 and Mtb.

## Results

### PWH and Mtb disease have higher HIV-1-specific neutralization breadth and potency

We first compared HIV-1 neutralization responses in ART-naive PWH who had confirmed Mtb disease (PWH/Active Mtb, n = 15) to PWH with no diagnosed or suspected active Mtb (PWH/No Mtb, n = 37, Fig. 1). Prior to any ART, all the PWH/Active Mtb samples were from individuals enrolled in a Mtb diagnostic trial in Uganda (20, 24). The PWH/No Mtb individuals were from the Mtb diagnostic trial (n = 16) and the AIDS Clinical Trial Group (ACTG) 5274 study (n = 21) (25). The PWH/No Mtb individuals had no confirmed or probable Mtb disease, although they were not evaluated for the presence of latent Mtb infection. The PWH/No Mtb as compared to PWH/Active Mtb (Table S1) and the Uganda as compared to ACTG 5274 (Table S2) individuals had significant demographic differences, such as the absolute CD4 T cell count. Thus, we used multivariate analyses to account for these differences.

**Figure 1.**
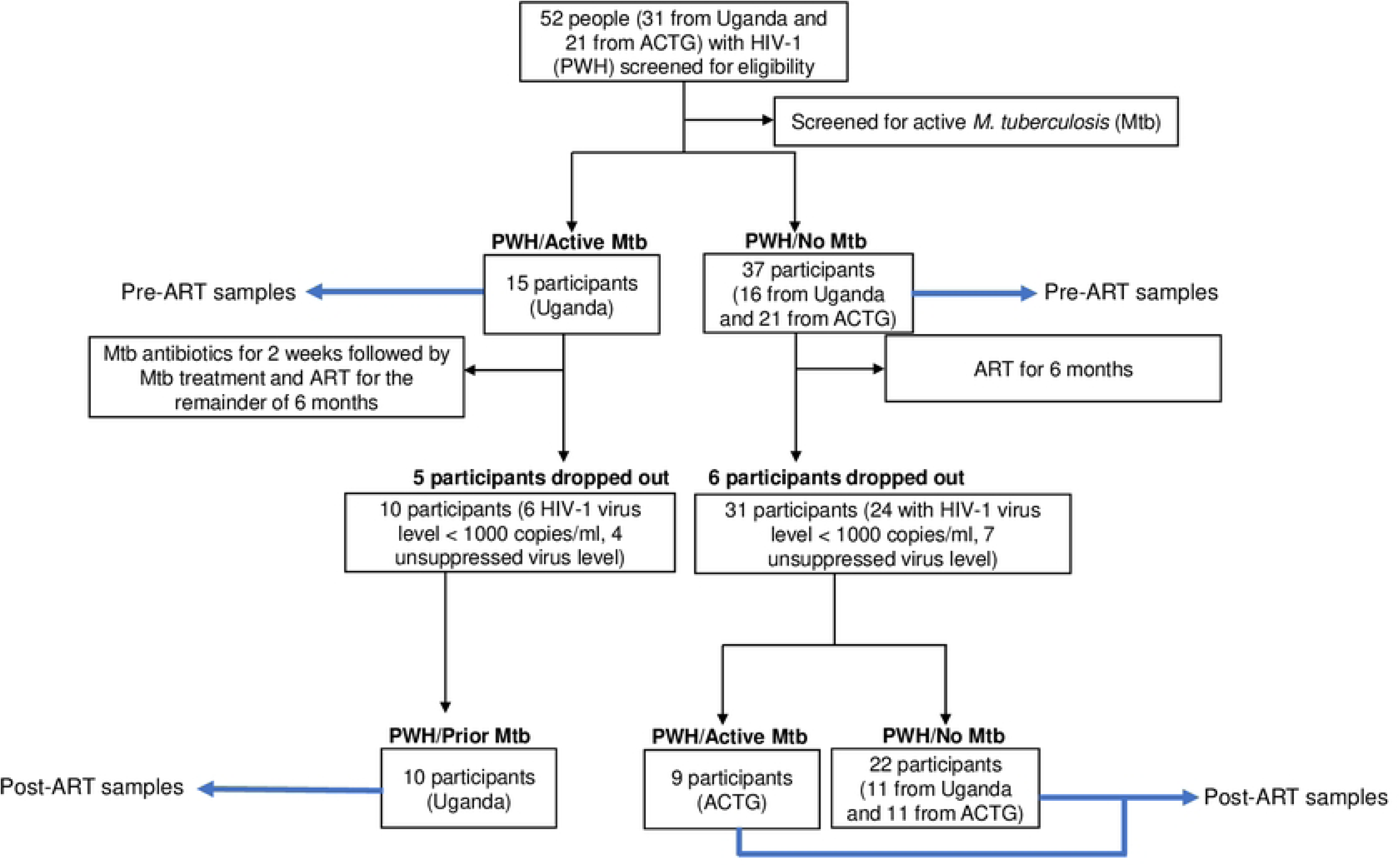
Participants and study design.

We examined responses against a panel of 12 different heterologous HIV-1 envelope glycoproteins (Envs) using the TZM-bl neutralization assay to compare HIV-1 neutralizing antibodies (nAbs) among the two groups (39). Neutralization against these 12 Envs recapitulates responses against over 200 globally isolated HIV-1 variants (27). HIV-1 neutralizing responses were compared using a breadth and potency (BP) score as described previously (38). Briefly, the neutralization BP score consists of an average of the log normalized percent neutralization against each virus at the highest tested plasma concentration (1:50 dilution). BP scores range from a minimum of 0 indicating no neutralization capacity to 1, which represents the ability to neutralize all twelve variants. Importantly, percent neutralization observed at the highest tested plasma dilution strongly correlates with both area under the neutralization curve and the plasma dilution required to achieve half-maximal inhibitory concentration (ID_50_) calculated using a series of diluted plasma (30, 31). Furthermore, the neutralization BP score strongly correlates with the neutralization breadth, defined as the percent of the global Env panel neutralized at greater than 50% at the highest tested plasma dilution (Fig. S1) (30, 31).

Prior to ART initiation, PWH/Active Mtb as compared to the PWH/No Mtb had significantly higher neutralization BP scores (p < 0.0001, Fig. 2A). In multi-variate linear regression analysis, PWH/Active Mtb as compared to PWH/No Mtb had around 0.26-unit higher BP score (95% confidence interval (CI) 0.15 – 0.37, p < 0.0001) after adjusting for gender, log_10_ plasma virus level, absolute CD4 count, and age (Table 1). Neutralization fingerprints were compared among the PWH/Active Mtb and PWH/No Mtb groups to highlight potential differences in antibody specificities among the groups. Prior to starting ART, heat map analysis showed that most plasma samples separated into three clusters with 100% bootstrap support (Fig. 2B). Cluster 3 contained only PWH/No Mtb samples, and these samples demonstrated less than 50% neutralization against nearly all Env variants except subtype A X398F1. Neutralization capacity against eight other Envs also differentiated PWH/Active Mtb and PWH/No Mtb (Fig. S2).

**Table 1.** Predictors of neutralization BP score prior to ART initiation.

| | Estimate ($\beta$) | 95% confidence interval | p-value |
| --- | --- | --- | --- |
| (Intercept) | 0.002 | -0.59 to 0.59 | 0.99 |
| Mtb | 0.26 | 0.15 to 0.37 | <0.0001 |
| Age | 0.001 | -0.004 to 0.006 | 0.62 |
| Male | -0.03 | -0.13 to 0.08 | 0.54 |
| CD4 count (cells / mm<sup>3</sup>) | 0.0003 | -0.0001 to 0.0007 | 0.16 |
| $\log_{10}$ plasma virus (copies / ml) | 0.05 | -0.05 to 0.14 | 0.33 |

**Figure 2.**
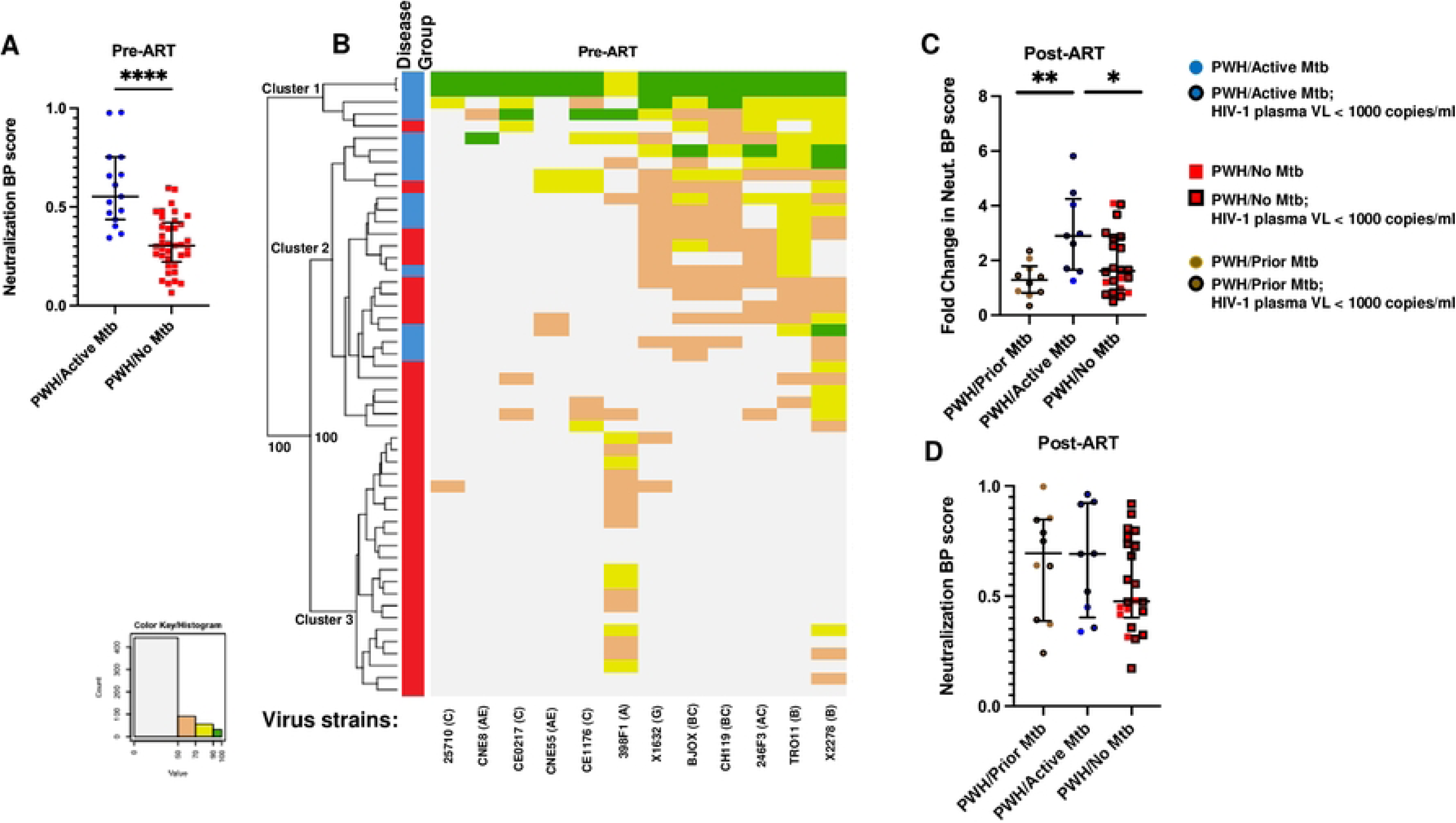
Mtb disease enhances the HIV-1 antibody response. (A & D) Neutralizing breadth potency (BP) score among (A) pre-ART and (D) post-ART plasmas. (B) Heatmap showing clustering of PWH/Active Mtb and PWH/No Mtb plasma samples before ART. Rows are individual plasma samples and columns are the 12 different Env variants. The left-hand side shows the clustering with bootstrap support. The bars to the left of the heatmap indicate either PWH/Active Mtb (blue) or PWH/No Mtb (red) plasma samples. The color histogram on the bottom left shows levels of neutralization: green 90 – 100%, lime green 70 – 90%, peach 50 – 70%, and light grey 0 – 50%. Below the heatmap, the column labels indicate the Env name and subtype in bracket. (C) Fold change of neutralization BP score at follow-up relative to baseline among post-ART plasmas. In all graphs, PWH/Treated Mtb (brown circles), PWH/Active Mtb (blue circles), and PWH/No Mtb (red squares). In (C & D), the brown, blue and red symbols with black boundaries represent those with post-ART plasma virus levels below 1000 copies per milliliter. Lines and error bars show median and interquartile ranges. Asterisks (*), (**), and (****) denote p-values < 0.05, <0.005, and less than 0.00005 respectively and are calculated based on an unpaired t-test with Welch’s correction (A) and Mann-Whitney test (D).

After enrollment, all participants initiated ART, and follow-up samples around six months later were available from 41 of the 52 individuals (Fig. 1). PWH/Active Mtb at baseline also started Mtb treatment based on national guidelines, and follow-up samples were collected after Mtb therapy completion (PWH/Prior Mtb, n = 10, Table S3). Nine of the PWH/No Mtb at entry had diagnosed or suspected Mtb disease a median of 140 (range 105 – 169) days after starting ART. These nine were either assigned to isoniazid prophylaxis (n = 4) or empiric Mtb treatment (n = 5) per ACTG 5274 protocol. These individuals were categorized as post-ART PWH/Active Mtb. None of the remaining PWH/No Mtb at baseline that were subsequently on ART alone (n = 11), or also on isoniazid prophylaxis (n = 3), or empiric Mtb treatment (n = 8) were diagnosed or suspected to have Mtb disease at follow-up (post-ART PWH/No Mtb). The post ART PWH/Active Mtb group had fewer men with lower absolute CD4 T cell counts compared to the other two groups although these differences were not statistically significant (Table S3). Post ART initiation, PWH/Active Mtb had a significantly higher ratio of follow-up to baseline neutralization BP score (median ratio 2.9, range 1.2 – 5.8) compared to PWH/Prior Mtb (median ratio 1.3, range 0.4 – 2.3) and PWH/No Mtb (median ratio 1.6, range 0.5 – 6.8) (Fig. 2C, p = 0.01, one-way ANOVA). In multivariate linear regression analysis, developing Mtb disease after starting ART associated with around 1.6-fold higher follow-up to baseline neutralization BP ratio (95% CI 1.0 – 2.6, p = 0.05) after accounting for absolute CD4 T cell count, plasma virus level, and gender. This suggests that active Mtb that develops after starting ART also associates with higher HIV-1 nAb responses. Furthermore, after ART initiation, neutralization BP score was relatively stable for PWH/Prior Mtb after finishing Mtb treatment and for the PWH/No Mtb individuals (Fig. 2D). In contrast to the ART naïve analysis, post-ART PWH/Active Mtb samples did not have significant separation in a neutralization heat map compared to the other two groups (Fig. S3).

### Mtb disease in ART-experienced PWH associates with a higher antibody dependent celluar cytotoxicity (ADCC)

Besides neutralization, antibodies can induce effector functions through Fc domains, such as natural killer (NK) cell-mediated ADCC. We and others have previously demonstrated that ADCC and neutralization responses are poorly correlated, which implies that they are independent functions (31,32,40,41). We examined ADCC against ten of the reference panel Envs among the groups before and after ART initiation using an assay that measures the killing of infected cells only and not of uninfected cells with bound shed Envs (32). An ADCC BP score was estimated in a similar manner as neutralization BP with 0 and 1 indicating no and 100% ADCC against all the strains in the panel respectively (40). Prior to starting ART, the ADCC BP score was not significantly different between the PWH/Active Mtb and PWH/No Mtb in both univariate (Fig. 3A) and multi-variate analysis (data not shown). After starting ART, PWH and active Mtb had a significantly higher ADCC BP score (p = 0.006, one-way ANOVA, Fig. 3B) and a higher ratio of follow-up to baseline ADCC BP (p = 0.0005, one-way ANOVA, Fig. 3C) compared to PWH/Prior Mtb and PWH/No Mtb. In multi-variable analysis, those with active Mtb disease after ART initiation had ADCC BP ratio around 0.18 units higher (95% CI 0.06 to 0.31, p = 0.004) after accounting for absolute CD4 count, gender, and plasma virus level (Table 2). In contrast to our prior observations, neutralization, and ADCC BP scores were moderately correlated prior to (r = 0.52, p < 0.0001, Fig. 3D, Pearson rank correlation) but not after starting ART (r = 0.28, p = 0.07, Fig. 3E). This pre-ART correlation was stronger in those with Mtb disease (r = 0.72, p = 0.003) as compared to the PWH/No Mtb (r = 0.32, p = 0.06). Post-ART, neutralization and ADCC BP scores showed a moderate statistical trend in correlation among those with active Mtb disease (r = 0.57, p = 0.10) and not those with prior or no Mtb (r = 0.23, p = 0.21). Thus, active Mtb disease associates with higher ADCC after but not before starting ART. Furthermore, neutralization and ADCC breadth and potency are primarily correlated among those with active Mtb.

**Table 2.** Predictors for the ratio of follow-up to baseline ADCC BP score.

| | Estimate ($\beta$) | 95% confidence interval | p-value |
| --- | --- | --- | --- |
| Intercept | 1.15 | 1.05 to 1.26 | <0.0001 |
| Active Mtb disease at follow up | 0.18 | 0.06 to 0.31 | 0.004 |
| Male | -0.014 | -0.24 to -0.04 | 0.006 |
| Post-ART CD4 + T cells (cells/mm<sup>3</sup>) | 0.0001 | -0.0001 to 0.0004 | 0.45 |
| Post-ART plasma virus level (copies/ml) | 0.002 | -0.03 to 0.03 | 0.87 |

**Figure 3.**
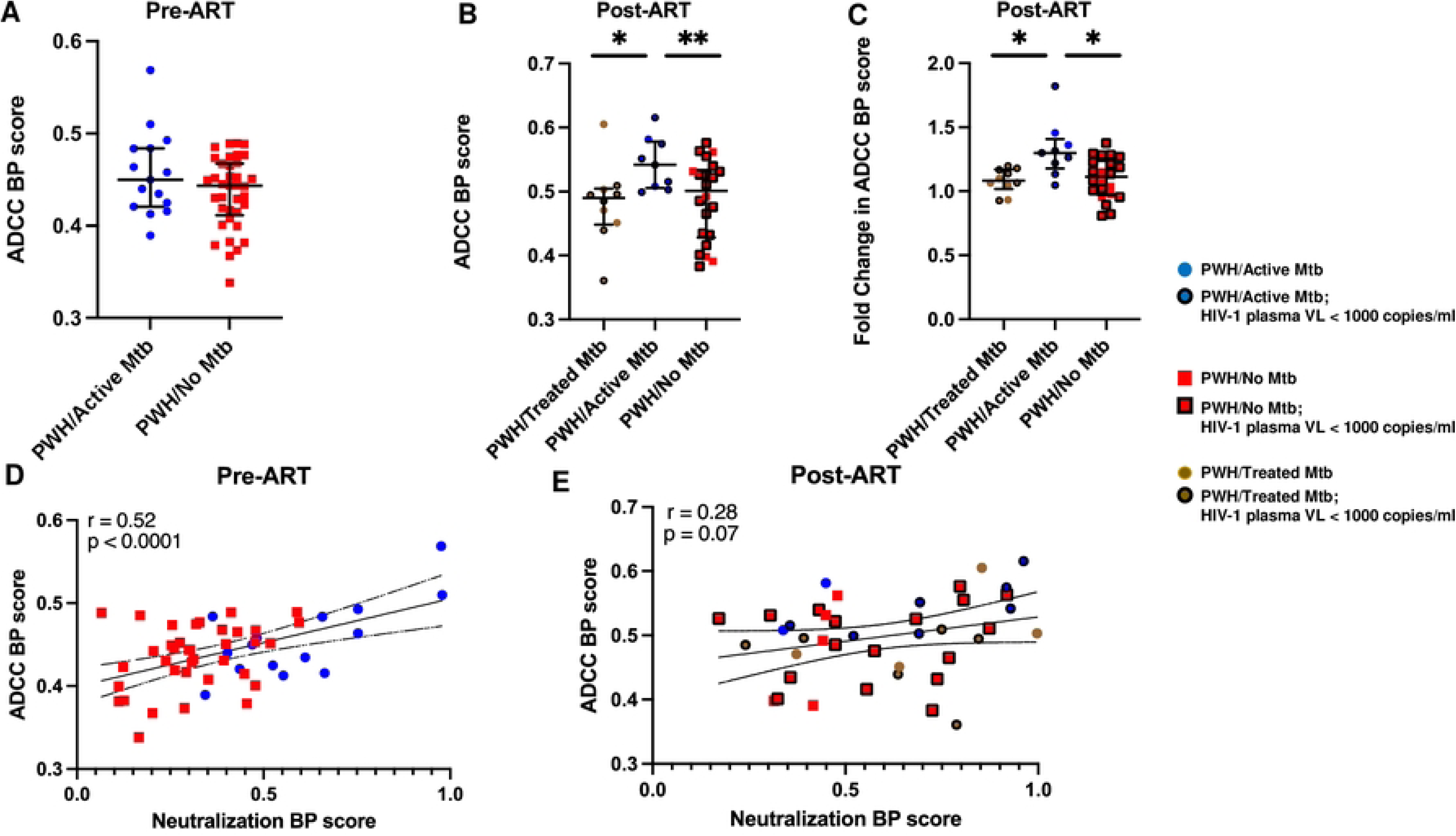
Plasma effector function activity associates with neutralizing activity. (A & B) ADCC breadth potency (BP) score among (A) pre-ART and (B) post-ART plasmas. (C) Fold change of ADCC BP score at follow-up relative to baseline. In (A – C) plasma from PWH/Treated Mtb (brown circles), PWH/Active Mtb (blue circles), and PWH/No Mtb (red squares). Lines and error bars show median and interquartile ranges. Asterisks (*) and (**) denote p-values < 0.05 and < 0.005 calculated based on an unpaired t-test with Welch’s correction. (D & E) Pearson’s correlation plots depict associations for (D) pre-ART ADCC BP score with neutralization BP score, and (E) post-ART ADCC BP score with neutralization BP score. In (B, C, & E), the symbols with boundaries represent those with post-ART plasma virus levels below 1000 copies per milliliter. Lines on correlation plots show the 95% confidence bands of linear regression.

### Factors associated with antibody breadth and potency

We next aimed to understand if the enhanced HIV-1 antibody responses observed in those with active Mtb disease were associated with previously identified factors deemed important for humoral breadth. Broad and potent HIV-1 antibody responses primarily develop because of a high level of virus replication, longer duration of infection, and exposure to a greater number of Env variants (42, 43). High plasma virus levels and low absolute CD4 counts serve as rough surrogates for greater virus replication and longer duration of HIV-1 infection respectively. Our previous multi-variable linear regression analysis showed that these factors did not independently associate with neutralization BP score after accounting for Mtb disease status (Table 1). Both longitudinal and cross-sectional studies have shown that Mtb disease increases HIV-1 replication (44, 45). Increased rounds of virus replication generate a larger number of genetic variants. Thus, Mtb disease may augment humoral responses by increasing the diversity of the HIV-1 Env antigen encountered by antibody-producing cells. We isolated a total of 341 SGA Env pre-ART sequences from a subset of the PWH/Active Mtb and PWH/No Mtb individuals. We did not analyze samples after ART initiation because ART distorts Env representation. Similar number of individuals and Envs per PWH were examined from PWH/Active Mtb and (n = 14, median Envs per subject 11, range 3 – 19) and PWH/No Mtb (n = 15, median Envs per subject 12, range 3 – 27, p = 0.32). Env sequences from different individuals clustered independently, although one PWH/No Mtb had sequences in more than one cluster suggesting a possible super-infection (46) (Fig. 4A). There was no significant difference in Env genetic diversity among the PWH/Active Mtb and PWH/No Mtb (Fig. 4B). Furthermore, Env genetic diversity did not correlate with pre-ART neutralization or ADCC BP scores (Fig. 4C and D). These analyses suggest that characteristics previously associated with HIV-1 antibody breadth and potency do not account for the enhanced humoral responses observed in PWH and Mtb disease.

**Figure 4.**
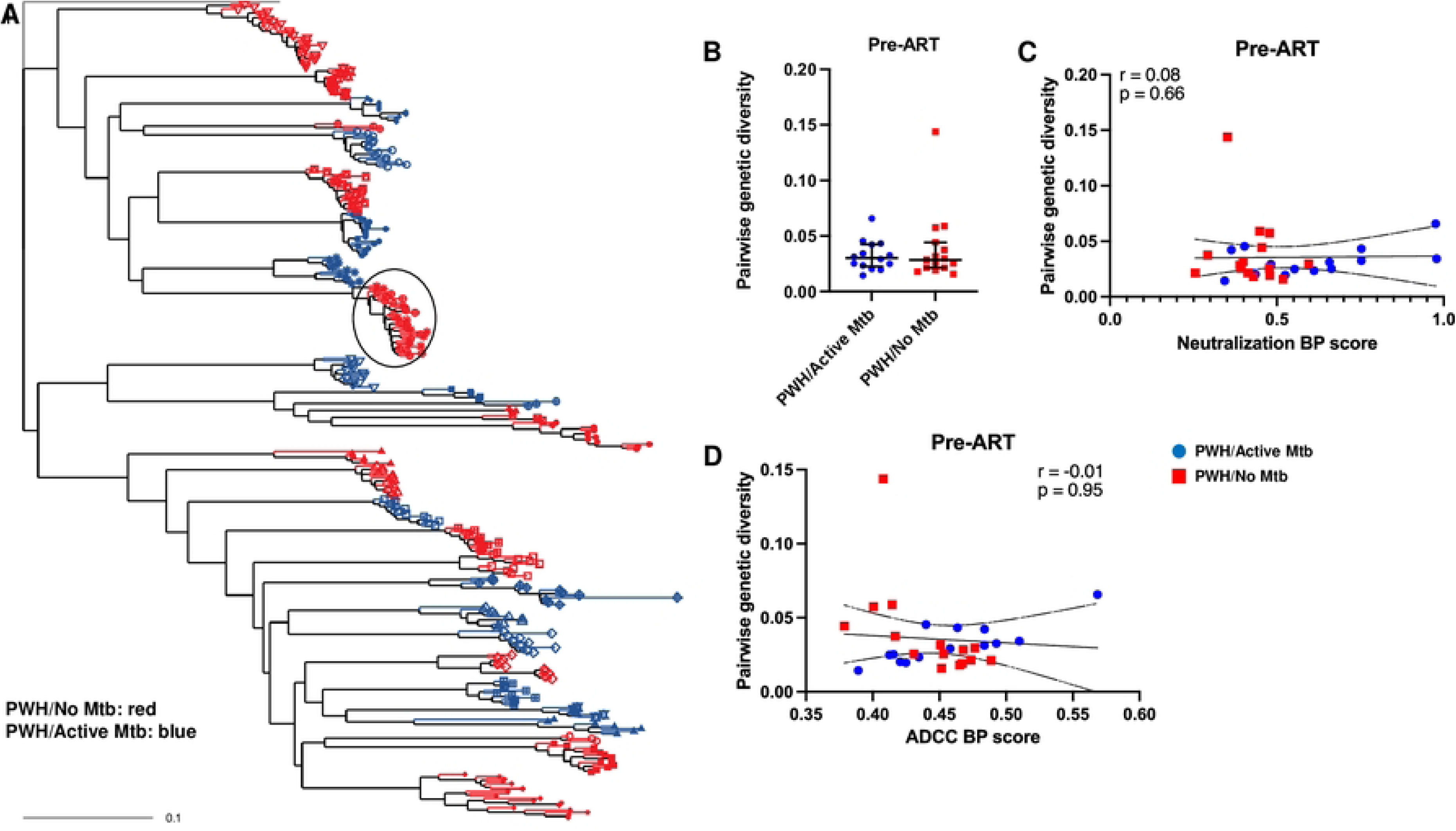
Viral Env diversity does not associate with neutralization and ADCC BP scores. (A) Maximum likelihood phylogenetic tree for PWH/Mtb (blue) and PWH/No Mtb (red). Different symbols at the tips indicate sequences from different individuals, and the circle shows interspersed sequences from two PWH/No Mtb. (B) Pairwise genetic diversity of PWH/Active Mtb (blue circles) as compared to PWH/No Mtb (red squares). Lines and error bars show median and interquartile ranges. (C-D) Correlation plots depict associations for (C) neutralization BP score and (D) ADCC BP score with pairwise genetic diversity in the PWH/Active Mtb (blue circles) and PWH/No Mtb (red squares). The r and p values indicate Spearman correlation statistic and lines depict linear regression with a 95% confidence interval.

### Env mutations are present at different frequencies

Antibodies that target conserved structures on the HIV-1 Env generally impart broad and potent nAb responses (47). Changes at specific amino acid sites that affect this conserved structure account for escape from bnAbs. We hypothesized that escape from the differential antibody pressure will yield HIV-1 Envs with unique amino acid signatures among PWH/Active Mtb as compared to PWH/No Mtb. We used the GenSig tool (https://www.hiv.lanl.gov/content/sequence/GENETICSIGNATURES/gs.html) to identify amino acids differentially present in the PWH/Active Mtb and PWH/No Mtb Env sequences. This analysis compares the proportion of each single amino acid in the PWH/Active Mtb and the PWH/No Mtb sequences with a phylogenetic correction to minimize false positives due to lineage effects (48). We used a false discovery rate (FDR) of Q < 0.1 to account for the multiple testing. Overall, there were thirty amino acids and eight predicted asparagine (N) linked glycosylation sites (PNGS) present at different frequencies in the PWH/Active Mtb and PWH/No Mtb sequences (Table S4 and S5). Among these differentially expressed motifs, thirteen sites have previously been determined to influence sensitivity to various HIV-1 bnAbs (Table 3). Four of these thirteen differentially present amino acids have been previously associated with both increased sensitivity and resistance depending on the Env context and antibody under consideration. PWH/Active Mtb and PWH/No Mtb sequences were enriched for seven and two of the remaining nine amino acid changes associated with resistance to bnAbs respectively (p = 0.06, Fisher’s exact test). Interestingly, the seven changes enriched in the PWH/Active Mtb sequences and associated with increased resistance to various bnAbs were all present in the Env surface unit (gp120). In contrast, the two resistance-inducing amino acids present at higher frequency in the PWH/No Mtb sequences were in the transmembrane membrane domain (gp41). Env variable loop length and the number of glycosylation sites also influence neutralization (30, 48). There was no significant difference in the variable loop lengths and the number of predicted glycosylation sites in the PWH/Active Mtb and PWH/No Mtb sequences (data not shown).

**Table 3:** Signature amino acids different in PWH/Active Mtb sequences and association with neutralization.

| AA at HXB2 position | AA association with neutralization | Odds ratio (test AA) (PWH/Active Mtb vs. PWH/No Mtb sequences) | p-value | Q-value |
| --- | --- | --- | --- | --- |
| T31 | T associated with resistance | 0 (K) | 0.006 | 0.05 |
| K33 | K associated with resistance | 0 (I) | 0.02 | 0.08 |
| D62 | E associated with resistance | Infinite (I) | 0.02 | 0.08 |
| T232 | T associated with sensitivity | Infinite (K) | 0.01 | 0.08 |
| V270 | V associated with resistance | Infinite (V) | 0.004 | 0.04 |
| V275 | K associated with resistance | 20 (K) | 0.01 | 0.09 |
| T278 | N eliminates a PNGS at site 276 and is associated with resistance and sensitivity | Infinite (N) | $2.9 \times 10^{-5}$ | $7.2 \times 10^{-5}$ |
| E293 | N associated with PNGS at 293 and with resistance | 3.6 (N) | 0.0006 | 0.002 |
| T297 | I and T associated with resistance and sensitivity | 0 (I) | 0.02 | 0.08 |
| N301 | Change from N eliminates a PNGS at site 301 associated with resistance and sensitivity | 2.7 (N) | 0.003 | 0.008 |
| N386 | Change from N eliminates a PNGS at site 386 associated with resistance and sensitivity | 11.4 (Y) | 0.005 | 0.01 |
| N750 | S associated with resistance | 0 (S) | 0.003 | 0.008 |
| R853 | R associated with sensitivity | Infinite (I) | 0.004 | 0.04 |

HXB2: reference sequence, amino acid: AA, arginine: R, asparagine: N, aspartic acid: D, glutamic acid: E, isoleucine: I, lysine: K, serine: S, threonine: T, valine: V, tyrosine: Y, Human Immunodeficiency Virus-1: HIV-1, and *Mycobacterium tuberculosis*: Mtb.
The odds ratio (OR) is based on Fisher's exact test. An OR < 1 indicates that odds of test AA are lower in PWH/Active Mtb as compared to PWH/No Mtb sequences. An OR > 1 denotes odds of test AA are higher in PWH/Active Mtb as compared to PWH/No Mtb sequences.
The p-value is from the phylogenetically corrected Fisher exact test.
Q-value is from false discovery rate multiple testing corrections.

We next used the bnAb-Resistance Predictor (bnAb-ReP), to estimate susceptibility to various bnAbs for the 341 different Envs (37). BnAb-Rep is an *in-silico* algorithm that uses machine learning to generate a resistance probability (ranging from 0 to 1) for different bnAbs based on Env sequences only. Envs are deemed resistant to a specific bnAb if they have a predicted probability below 0.5. Importantly, this algorithm has high prediction accuracy relative to *in-vitro* results. None of the 33 bnAbs assessed was predicted to neutralize all 341 Envs (data not shown). We focused on the few bnAbs targeting different Env epitopes that were both predicted to neutralize the majority of the 341 different Envs and also had a previously published prediction performance area under the curve above 0.85 (Table 4 and S6) (37). Envs from PWH/Active Mtb as compared to PWH/No Mtb had more than a 1.5 greater chance of predicted resistance to the CD4 binding site and V1-V2 directed bnAbs (Table 4). In contrast, PWH/Active Mtb as compared to the PWH/No Mtb Envs were predicted to have around a 50% or lower chance of being resistant to the V3 loop and membrane-proximal external region (MPER) directed bnAbs. This differential enrichment for the sequence motifs associated with bnAb resistance among PWH/Active Mtb and PWH/No Mtb further suggests unique antibody selection pressure in the two groups.

**Table 4.**
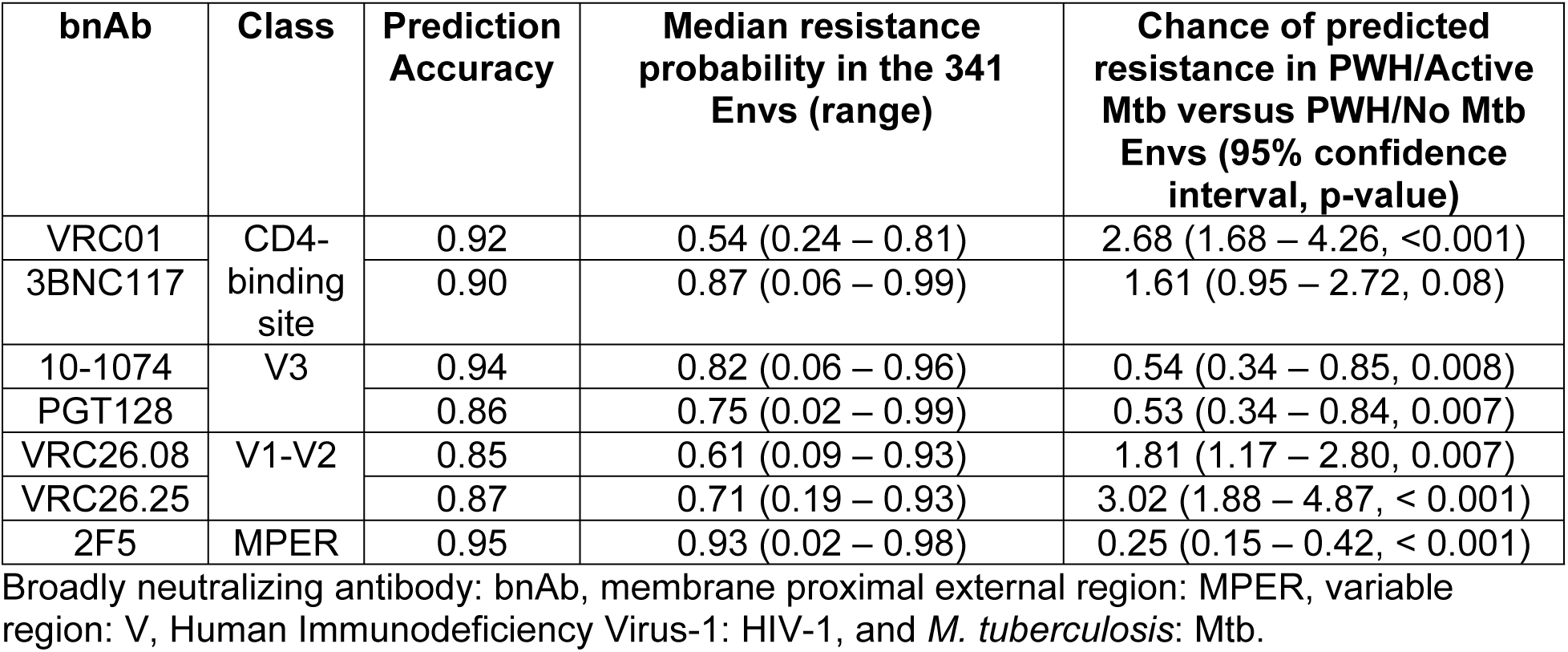
Predicted bnAb susceptibility among PWH/Active Mtb as compared to PWH/No Mtb.

| bnAb | Class | Prediction Accuracy | Median resistance probability in the 341 Envs (range) | Chance of predicted resistance in PWH/Active Mtb versus PWH/No Mtb Envs (95% confidence interval, p-value) |
| --- | --- | --- | --- | --- |
| VRC01 | CD4-binding site | 0.92 | 0.54 (0.24 – 0.81) | 2.68 (1.68 – 4.26, <0.001) |
| 3BNC117 |  | 0.90 | 0.87 (0.06 – 0.99) | 1.61 (0.95 – 2.72, 0.08) |
| 10-1074 | V3 | 0.94 | 0.82 (0.06 – 0.96) | 0.54 (0.34 – 0.85, 0.008) |
| PGT128 |  | 0.86 | 0.75 (0.02 – 0.99) | 0.53 (0.34 – 0.84, 0.007) |
| VRC26.08 | V1-V2 | 0.85 | 0.61 (0.09 – 0.93) | 1.81 (1.17 – 2.80, 0.007) |
| VRC26.25 |  | 0.87 | 0.71 (0.19 – 0.93) | 3.02 (1.88 – 4.87, < 0.001) |
| 2F5 | MPER | 0.95 | 0.93 (0.02 – 0.98) | 0.25 (0.15 – 0.42, < 0.001) |

Broadly neutralizing antibody: bnAb, membrane proximal external region: MPER, variable region: V, Human Immunodeficiency Virus-1: HIV-1, and *M. tuberculosis*: Mtb.

### All or non-HIV-1 antibodies not elevated in Mtb disease

We hypothesized that Mtb disease does not enhance the production of all antibodies, but rather it is HIV-1 specific in PWH. First, we examined total antibody levels to assess if Mtb disease associates with a generalized antibody increase. Total IgG levels were not significantly higher among those with active Mtb both before and after starting ART (Fig. 5A and B). Furthermore, different immunoglobulin isotypes were not significantly different among those with and without Mtb disease both before and after ART (Fig. S4A - L). In prior studies, HIV-1 specific neutralizing breadth and potency has also been associated with elevated total immunoglobulin (IgG) levels (49), but there was no correlation between total IgG levels and neutralization BP score, both before and after ART (Fig. S5A and B). In previous studies from our group and others, a higher IgG/IgA ratio associates with a greater ADCC (31,40,50–52). Similarly, in multi-variable linear regression analysis, the pre-ART IgG/IgA ratio positively correlated with ADCC after accounting for baseline demographics (Table S7).

**Figure 5.**
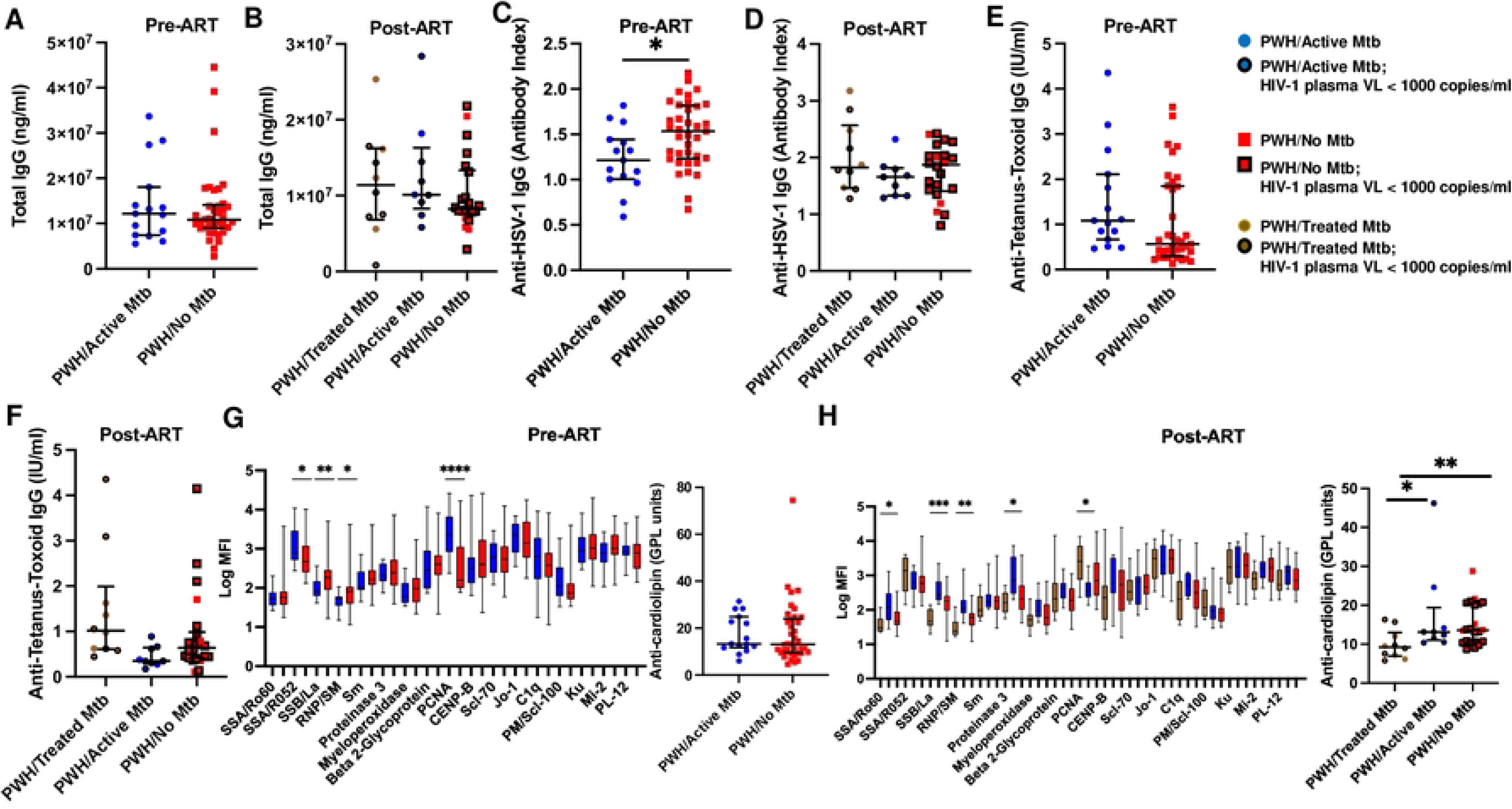
Mtb disease does not affect the levels of immunoglobulins and non-HIV-1 antibodies. (A - F) Concentrations of total IgG, anti-HSV-1 IgG, and tetanus-toxoid specific IgG (A, C & E) before and (B, D & F) after ART. Lines and error bars show median and interquartile ranges. (G & H) Mean fluorescent intensity (MFI) of autoantibodies and phospholipid IgG (GPL) levels of anticardiolipin (G) before and (H) after ART. In (G & H), lines and whiskers show median and minimum to maximum range respectively. Graphs show PWH/Active Mtb (blue circles), PWH/No Mtb (red squares), and PWH/Treated Mtb (brown circles), and the symbols with black boundaries represent those with post-ART plasma virus levels with less than 1000 HIV-1 copies/ml. Asterisks (*), (**), (***), (****) denote p-values < 0.05, < 0.005, < 0.0005, and < 0.00005 calculated based on an unpaired t-test with Welch’s correction.

We next examined levels of antibodies that likely preexist in PWH prior to the development of Mtb disease. Antibodies against tetanus and HSV-1/2 are commonly detected in HIV-1 seropositive adults (53, 54). In contrast to the HIV-1 humoral responses, PWH/Active Mtb had lower anti-HSV-1 IgG titers prior to starting ART although this was not statistically significant after accounting for demographic differences in multi-variable analysis (Fig. 5C and data not shown). There was no difference in anti-HSV-1 IgG after starting ART among the three groups (Fig. 5D). Anti-tetanus IgG titers were similar in PWH/Active Mtb and PWH/No Mtb prior to ART (Fig. 5E). The PWH/Prior Mtb had higher anti-tetanus IgG compared to those with active Mtb after ART, but this was not statistically significant in multi-variate analysis. (Fig. 5F and data not shown).

Finally, we assessed the levels of antibodies that are potentially generated de-novo during acute disease episodes. Numerous infectious diseases have been associated with the emergence of autoantibodies (55, 56). We estimated autoantibody levels using Luminex-based mean fluorescence intensity (MFI) against beads coated with seventeen different autoantigens and ELISA to measure anti-cardiolipin phospholipid IgG levels. Before and after ART, some autoantibody levels were different among the groups in univariate analysis (Fig. 5G and H), but no group was consistently higher or lower. The presence of Mtb disease also did not have a significant association with any of the assessed autoantibody levels after accounting for multiple comparisons. None of the autoantibodies associated with either neutralization or ADCC BP score before or after ART in multi-variable linear regression analysis after accounting for multiple comparisons (data not shown).

### Mtb infection does not elicit cross-reactive responses

Mtb disease association with enhanced HIV-1 neutralizing response in the absence of a generalized increase in antibodies may result from cross-reactivity. A previous study suggested that Mtb and HIV-1 share some epitope similarities (57). We assessed HIV-1 plasma neutralization activity in 32 HIV-1 seronegative participants, some with a previous history of Mtb infection to examine possible cross-reactivity. Of these 32 participants, eight were healthy control, sixteen had latent Mtb, four had active Mtb disease, and four had just finished treatment for active Mtb (Table S8) (26). No individual’s plasma demonstrated greater than 50% inhibition against any of the five tested reference panel Envs (246F3, BJOX, CE1176, 25710, and CNE8) (Fig. 6). Thus, Mtb infection and disease do not elicit nAbs that cross-react with HIV-1 Envs.

**Figure 6.**
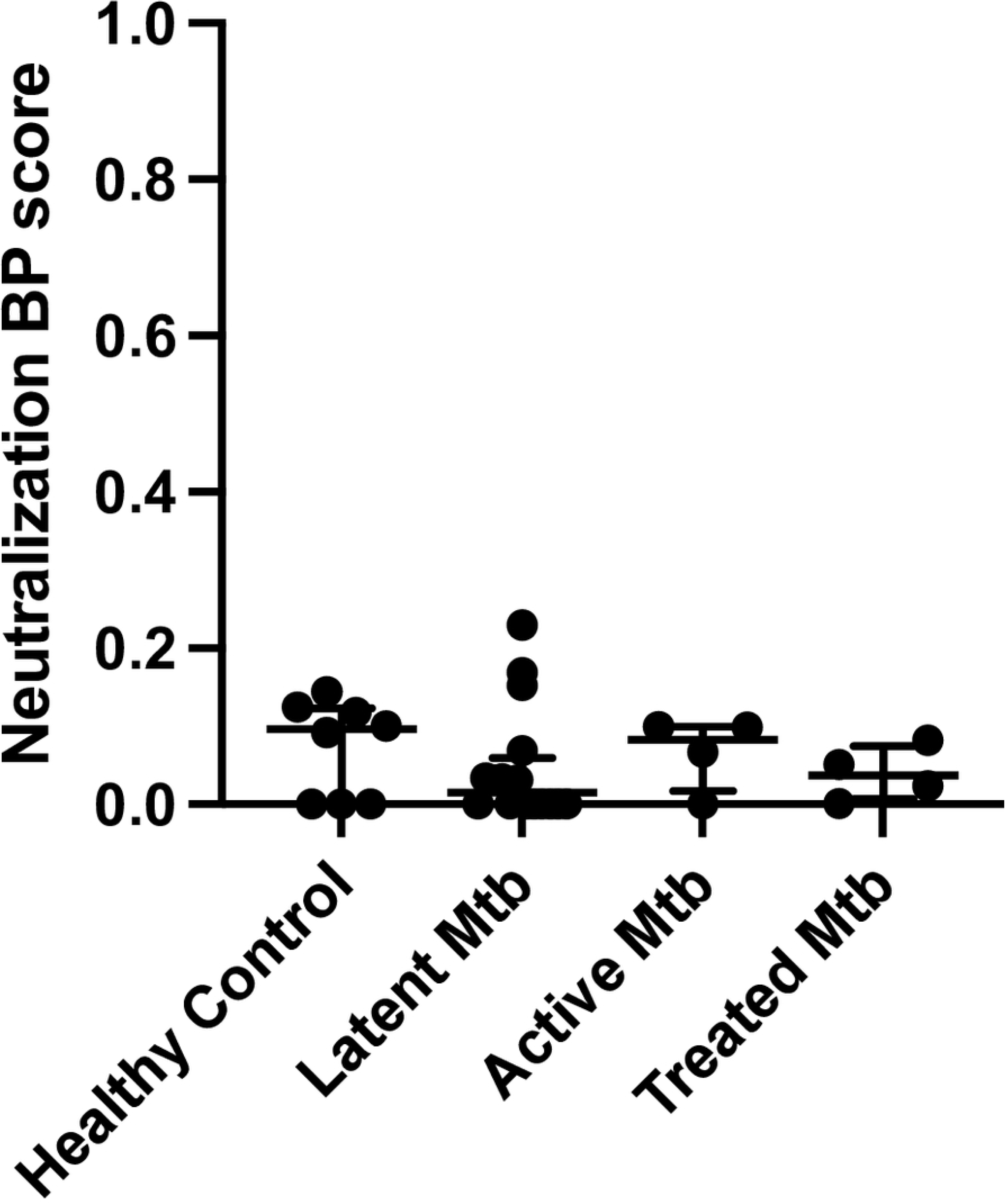
Mtb does not induce cross-reactive HIV-1 antibodies. Neutralization BP score derived from plasma neutralization against five HIV-1 isolates among those who have no Mtb (healthy controls (*n* = 8), latent Mtb (*n* = 16), active Mtb (*n* = 4) and treated Mtb (*n* = 4). Lines and error bars show median and interquartile ranges.

### Active Mtb disease is associated with a unique inflammatory profile

In the absence of cross-reactivity and a generalized antibody increase, we hypothesized that Mtb-mediated systemic inflammation may enhance HIV-1 antibody response as a bystander effect. Both the development of broad and potent neutralizing activity and Mtb disease have been associated with immune activation (58, 59). We measured a diverse set of analytes deemed important for lymph node germinal center (GC) formation, B cell development, and antibody production using a human B cell multiplex panel in the pre-ART plasmas. Only seven (interleukin-6 (IL-6), C–X–C motif chemokine 10 (CXCL10), chemokine motif ligand 5 (CCL5, RANTES), IL-10, a proliferation-inducing ligand (APRIL), B-cell activating factor (BAFF), and soluble CD40 ligand (sCD40L)) of the fifteen measured analytes deemed important for B cells and antibodies yielded values above the limit of detection (LOD) for the majority of samples. Prior to any treatment, IL-6, APRIL, and BAFF were significantly higher in PWH/Active Mtb as compared to PWH/No Mtb (Fig. 7A - C), but only the IL-6 difference was statistically significant after accounting for multiple comparisons. Log_10_ IL-6 level (β = 0.12, 95% CI 0.04 –0.22, p = 0.007) and Log_10_ BAFF level (β = 0.11, 95% CI 0.02 – 0.21, p = 0.01) also predicted neutralization BP score after accounting for age, gender, absolute CD4 count, and log_10_ plasma virus level in multi-variable linear regression analysis. As expected, IL-6 and BAFF did not predict pre-ART neutralization BP score if Mtb disease status was included in a multi-variable linear regression model (Table 5), because IL-6 (Fig. 7A) and BAFF (Fig. 7C) levels associate with Mtb disease. Thus, IL-6, BAFF, and Mtb disease status are inter-dependent.

**Table 5.** Predictors of neutralization BP score prior to ART initiation including plasma cytokines.

| | Estimate ($\beta$) | 95% confidence interval | p-value |
| --- | --- | --- | --- |
| (Intercept) | 0.037 | -0.62 to 0.69 | 0.91 |
| Mtb | 0.23 | 0.08 to 0.37 | 0.002 |
| Age | 0.001 | -0.004 to 0.006 | 0.72 |
| Male | -0.02 | -0.12 to 0.08 | 0.67 |
| CD4 count (cells / $\text{mm}^3$) | 0.0002 | -0.0001 to 0.0007 | 0.21 |
| $\text{Log}_{10}$ plasma virus (copies / ml) | 0.03 | -0.08 to 0.13 | 0.41 |
| $\text{Log}_{10}$ plasma IL-6 (pg/ml) | 0.042 | -0.06 to 0.14 | 0.41 |
| $\text{Log}_{10}$ plasma BAFF (pg/ml) | 0.0009 | -0.10 to 0.10 | 0.99 |

**Figure 7.**
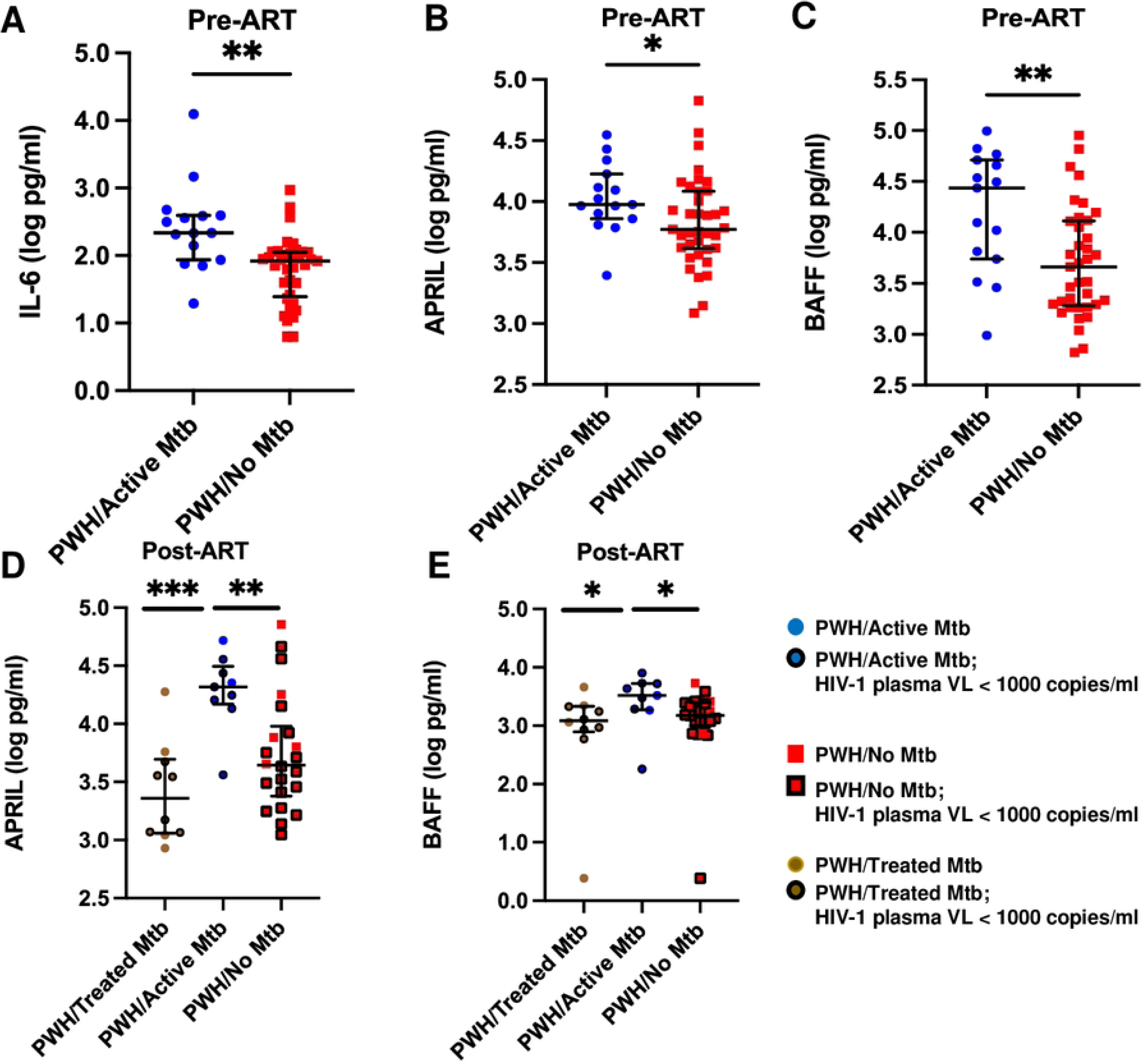
Plasma markers of germinal center activity associates with neutralizing activity. (A - C) Pre-ART (A – C) and post-ART concentrations (D – E) of IL-6 (A), APRIL (B & D), and BAFF (C & E) for PWH/Active Mtb (blue circles) PWH/No Mtb (red squares), and PWH/Treated Mtb (brown circles). In (D & E), the symbols with boundaries represent those with post-ART plasma virus levels below 1000 copies per milliliter. Lines and error bars show median and interquartile ranges. Asterisks (*), (**), and (***) denote p-values < 0.05, <0.005, and <0.0005 respectively and are calculated based on an unpaired t-test with Welch’s correction (A – D) and Mann-Whitney test (E).

After starting ART, APRIL (p = 0.0006, one-way ANOVA, Fig. 7D) and BAFF (p = 0.03, Kruskal-Wallis test, Fig. 7E) were significantly higher in PWH/Active Mtb as compared to the other two groups. Similar to our previous observations after ART initiation, the majority of individuals had plasma IL-6 levels below the LOD preventing us from doing IL-6 comparisons (20). Log_10_ APRIL levels (β = 1.0, 95% CI 0.2 – 1.9, p = 0.02) but not BAFF levels significantly associated with the post-ART to pre-ART neutralization BP ratio after adjusting for gender, plasma virus level, and post-ART CD4 count. Similar to the pre-ART analysis as expected, APRIL did not predict the follow-up to baseline neutralization BP ratio if Mtb disease status was included in a multi-variable linear regression model (data not shown), because APRIL and presence of Mtb disease are inter-dependent (Fig. 7D). This implies that Mtb disease-associates with higher IL-6, BAFF, and APRIL levels and this increase subsequently predicts HIV-1 neutralization breadth and potency.

## Discussion

The existence of mycobacteria-mediated heterologous protection against different pathogens has been extensively documented in humans (5,7,60). Despite the high burden of HIV-1 and Mtb co-infection, the impact of Mtb infection and disease on the humoral response against HIV-1 has not been investigated. Our study suggests that active Mtb disease associates with the emergence of a broad and more potent anti-HIV-1 neutralization response. After ART initiation, Mtb disease associates with both broader and more potent nAbs and ADCC. Moreover, greater antigen exposure, higher level of virus replication, longer duration of HIV-1 infection, generalized antibody increase, and cross-reactivity does not account for the augmented antibody responses in PWH/Active Mtb. Rather from our data, we speculate that Mtb disease alters cytokines important for antibody production in lymphoid tissues that harbor both pathogens, and this accounts for enhanced HIV-1 humoral immunity. Although broad and potent HIV-1 humoral responses do not provide virologic control or improve disease outcomes in ART naïve PWH (43,61,62), our observations have implications both for using antibodies as therapeutics and preventing HIV-1 acquisition and for developing strategies to induce optimal immunity in PWH.

We provide multiple independent lines of evidence that Mtb disease associates with enhanced HIV-1 nAbs. First, we observed that PWH/Active Mtb had around 0.26-unit higher neutralization BP score as compared to the PWH/No Mtb prior to ART initiation. This increase is similar to or greater than the difference between some second as compared to first-generation HIV-1 bnAbs (38). In general, the second as compared to first-generation bnAbs are more potent and can inhibit a larger diversity of HIV-1. Thus, the observed difference in neutralization capacity among PWH/Active Mtb as compared to PWH/No Mtb group is biologically meaningful. Second, we show that after starting ART, PWH and active Mtb disease have a greater relative increase in neutralization breadth and potency compared to PWH and prior Mtb or no Mtb disease. This suggests that developing Mtb disease after ART also associates with greater neutralization capacity. Third, the Mtb disease impact on HIV-1 humoral immunity is not transient, because although antibodies can increase soon after ART initiation (63), the PWH with either prior treated Mtb or no Mtb had relatively stable humoral responses over time. Notably, the PWH and no Mtb had the lowest antibody responses both prior to and after ART. Fourth, our sequence analysis suggests PWH/Active Mtb have differential enrichment for sequence motifs associated with bnAb resistance. In general, HIV-1 variants evolve to escape nAbs (30,64,65), and the presence of differentially susceptible strains implies that the virus population in individuals with Mtb disease is under unique antibody selection pressures. In aggregate, active Mtb disease associates with higher HIV-1 antibody response both prior to and after ART initiation.

Not all individuals with chronic HIV-1 infection develop broad and potent neutralization responses and the reasons that only a minority develop bnAbs remain uncertain (42, 66). Previous studies have shown that factors important for bnAb development include exposure to higher amounts and longer duration of diverse viruses. The Mtb-associated neutralization increase did not correlate with plasma virus level, absolute CD4 count, or envelope genetic diversity, which are surrogate markers for the antigen levels, duration of infection, and epitope variation respectively. In the absence of HIV-1, individuals with latent, active, or treated Mtb also did not have demonstrable HIV-1 antibody responses, and thus the observed results are not due to cross-reactivity. Three independent lines of evidence also imply that the increase in HIV neutralization capacity in the presence of Mtb disease does not merely reflect a generalized increase in antibody production. First, although chronic HIV-1 infection is associated with hypergammaglobulinemia and derangements in the IgG isotype proportions (67, 68), in general, there was no difference in total IgG, IgA, or IgM levels in the PWH/Active Mtb and PWH/No Mtb groups. Second, there was no difference in antibody levels that most likely pre-existed prior to Mtb disease, such as those against tetanus and HSV-1/2. Third, antibodies that target different autoantigens, which are often generated de-novo after infection, were neither consistently higher nor lower in the groups. In aggregate, this implies Mtb infection specifically enhances HIV-1 antibodies. We assessed antibody levels for the non HIV-1 antibodies, while we quantified antibody functionality (neutralization and ADCC) for the HIV-1 responses. HIV-1 nAb breadth and potency correlates with anti-Env IgG binding titers (49), and thus we would predict that total HIV-1 anti-Env antibody levels are also higher in the PWH and active Mtb disease. We concentrated on neutralization and ADCC because these are biologically more relevant than binding titers. In summary, the usual factors associated with higher and broader HIV-1 nAbs do not account for the Mtb linked enhanced HIV-1 antibody response.

We speculate that there are likely novel mechanisms for the Mtb-associated increase in HIV-1 antibody breadth and potency. During Mtb disease and chronic HIV-1 infection, both microorganisms can be found in lymph nodes, which are the primary site for B cell development and antibody production. Furthermore, both organisms may be present in granulomas formed during Mtb disease (22, 23), and granulomas are often classified as secondary lymphoid structures (69, 70). Mtb is known to engage pathogen recognition receptors (PRR), such as toll-like receptors (TLRs) and nucleotide-binding oligomerization domain-containing protein 2 (NOD2), and these are present on the lymphoid tissue-resident follicular dendritic cells (FDCs), CD4+ T follicular helper (TFH), and B cells (71). Mtb engagement of these receptors may influence antigen presentation and protein production by the adjacent HIV-1-laden FDCs and TFH (72). This may increase the amount of HIV-1 Env sampled by B cells in the lymph nodes that harbor both HIV-1 and Mtb. Longer and greater antigen presentation is important for the generation of HIV-1 bnAbs (73). Furthermore, Mtb disease likely increases tissue levels of soluble factors important for antibody production, such as IL-6, BAFF, and APRIL as indicated by our plasma analysis. Elevated IL-6 levels may promote B – TFH cell interactions, which is important for somatic hypermutation (SHM) (21, 74). The majority of HIV-1 bnAbs display extensive SHM (75), and Mtb-mediated IL-6 elevation may promote SHM among the developing HIV-1 antibodies in the lymphoid tissue that contain both HIV-1 and Mtb. Furthermore, elevated levels and correlations observed with BAFF/APRIL suggest that Mtb may help promote the survival and maintenance of B cells, especially some that may be autoreactive (76). The BAFF/APRIL cascade has been associated with the presence of autoimmune antibodies, and self-reactivity is deemed important in bnAb emergence (76). The multi-variate linear regression analysis suggests that these cytokines may drive neutralization breadth and potency after they are induced by Mtb disease. This possible model argues that HIV-1 antibody response may be enhanced in a bystander fashion in Mtb co-infected lymphoid tissues. Examining this potential biological mechanism will require the development of a co-infection model or lymphoid tissue from PWH and Mtb disease.

Similar to neutralization, ADCC breadth and potency was also higher in those with active Mtb disease, albeit only after starting ART. In addition, ADCC moderately correlated with nAb response, primarily in those with Mtb. ADCC requires antibody binding (Fab) portion attaching to the antigen and the antibody crystallizable (Fc) fragment engaging with Fc receptors (FcRs) on effector cells. Afucosylation and galactosylation in the antibody Fc domains are known to impact ADCC (77, 78). Mtb disease may influence both antibody binding to HIV-1 Env on infected cells and antibody Fc compositions. While broad and potent HIV-1 neutralization responses are mediated via bnAbs rather than a panoply of different antibodies (79), the antibody characteristics responsible for ADCC breadth and potency have not been defined. ADCC breadth and potency could potentially result from antibodies that target conserved Env epitopes, similar to bnAbs, along with or primarily because of unique Fc characteristics. Isolating and characterizing individual monoclonal antibodies from PWH and Mtb disease will provide further insights. We did not have PBMC to isolate HIV-1 specific memory B cells and characterize whether the nAbs and the antibodies that mediate cellular cytotoxicity target the same or different Env epitopes.

The inclusion of a relatively large number of individuals from multiple cohorts representing diverse geographical sites is a strength of the study. This study, however, is limited by the lack of PBMCs and tissue samples, and participant dropout at follow-up. Furthermore, in this investigation, we cannot decipher if the HIV-1 infection preceded active Mtb or vice versa. Although the order of infections is not known, our longitudinal observations suggest that Mtb disease development with prior documented HIV-1 infection is associated with HIV-1 nAb and ADCC enhancement. It remains uncertain if latent Mtb infection in the absence of active disease is also associated with similar changes in neutralization capacity and ADCC. It could be speculated that PWH and latent Mtb infection potentially also have more potent immune responses because PWH and latent Mtb have lower set point HIV-1 plasma virus levels (12). In general, however, antibodies do not have a significant impact on the amount of virus in the blood.

Previous studies have demonstrated that broad and potent nAb responses do not impact HIV-1 disease progression among individuals who do not have suppressed virus levels (43,61,62). Neutralization escape variants emerge quickly with the ongoing virus replication, and antibodies are ineffective against neutralization-resistant strains (65). Thus, our observations have implications for using HIV-1 antibodies as therapy and preventing acquisition. PWH with prior or current Mtb disease are more likely to have HIV-1 neutralization breadth and harbor bnAb resistant strains. Thus, future bnAb use to suppress viremia and prevent transmission will likely be less effective among HIV-1 infected populations with high prevalence of Mtb disease (80–83).

It is well known that Mtb disease worsens morbidity and mortality in PWH even if they are on suppressive ART (16). In addition, BCG vaccination is strictly contraindicated in HIV-1 infected children regardless of disease stage or treatment. Thus, mycobacterial infections cannot be used to enhance HIV-1 immune responses. Our results, however, could provide the scientific basis for future investigations aimed at understanding how Mtb disease may yield broad and potent HIV humoral responses, besides the usual unmodifiable factors, namely prolonged duration of infection and high plasma virus level (43, 62). Extrapolation of our observations would suggest that individuals with suppressed virus levels who undergo an intervention that mimics aspects of active Mtb disease may develop a broad and potent HIV-1 nAb and ADCC response. These augmented nAbs and ADCC could change the size and nature of the residual HIV-1 latent reservoir if they are effective against autologous strains. Examination of larger cohorts, quantifying ADCC against the autologous strains, and assessing correlations with measures of HIV-1 latency will be required to understand these issues in detail. Similar strategies are being pursued with bnAb infusion treatments (80). In contrast, novel insights gleaned from understanding the mechanism for Mtb antibody enhancement could be more practical for the majority of HIV-1 infected individuals worldwide.

## Materials and Methods

### Study participants

Samples were obtained from three different cohorts. The first set of samples were from consecutively enrolled individuals newly diagnosed with HIV-1 infection who were part of a Mtb diagnostic study in Kampala, Uganda as described previously (20, 24). Briefly, the presence and absence of Mtb disease was assessed with GeneXpert MTB/RIF test and two Mtb liquid and one solid culture on two expectorated sputum samples. PWH who had confirmed Mtb disease (PWH/Active Mtb, n = 15) had a positive GeneXpert MTB/RIF result and at least one sputum sample culture positive for the presence of Mtb. PWH with no diagnosed or suspected active Mtb (PWH/No Mtb, n = 16) had negative results with all Mtb tests. PWH/Active Mtb initiated standard six-month anti-Mtb therapy immediately, and ART was started approximately two weeks later. PWH/No Mtb began ART after HIV-1 diagnosis. All treatments were based on national and World Health Organization guidelines. Blood samples were also obtained from participants that returned for a follow-up visit around six months after the initial enrollment visit. At this point, none of the PWH/Active Mtb had ongoing TB disease, and they were classified as PWH/Prior Mtb (n = 10). After starting ART, none of the PWH/No Mtb had evidence of active Mtb (n = 11).

The second set of samples was from outpatient research clinics in South Africa, Haiti, Kenya, and India that were part of the ACTG 5274 trial (NCT01380080), which compared treatment strategies for preventing Mtb associated mortality in PWH (25). At enrollment and prior to ART, none of the PWH in the ACTG 5274 trial had confirmed or probable Mtb disease (n = 21). After starting ART, individuals were randomized to isoniazid prophylaxis or empiric Mtb treatment. Follow-up samples were obtained around six months after enrollment. At follow up, nine individuals were diagnosed with or presumed to have Mtb disease based on national standards and testing, including acid-fast bacilli smears, chest radiography, ultrasound, mycobacterial culture, and GeneXpert. Eleven individuals did not have confirmed or probable Mtb at the follow-up visit. In both the Uganda and A5274 cohort, none of the PWH/No Mtb were evaluated for the presence of latent Mtb at any time.

The final set of samples was from 32 HIV-1 uninfected individuals who either had diagnosed latent Mtb (n= 16), active Mtb disease (n = 4), recovered from active Mtb after treatment (n = 4), or no detectable Mtb exposure (n = 8) as described previously (26).

### Virus stocks and cell lines

Twelve reference Envs were obtained from the NIH AIDS reagent program (27). Of the twelve, ten reference Envs (TRO11, X2278, X1632, 246F3, BJOX, CE1176, 25710, 398F1, CNE8, and CNE55) were incorporated into replication-competent viruses (28, 29), and two reference Envs (CH1119 and CE0217) were pseudotyped using protocols detailed previously (30). All viral stocks generated were stored at −80°C and titers were determined on TZM-bl cells.

TZM-bl and HEK293T cells were obtained from the NIH AIDS reagent program. NK (CD16+KHYG-1) and MT4-CCR5-Luc cells were obtained and propagated as described previously (31, 32). TZM-bl and HEK293T cells were maintained in complete Dulbecco’s modified Eagle medium (DMEM) containing 10% fetal bovine serum (FBS) (Invitrogen), 2 mM L-glutamine (Invitrogen), 100 U/mL of penicillin, and 100 μg/mL of streptomycin. MT4-CCR5-Luc cells were maintained in complete Roswell Park Memorial Institute (RPMI) medium (Invitrogen) containing 10% FBS (Invitrogen), 2 mM L-glutamine (Invitrogen), 100U/mL of penicillin, and 100 μg/mL of streptomycin. NK (CD16+KHYG-1) cells were sustained in complete RPMI supplemented with 0.1 mg/ml Primocin (InvivoGen), 25 mM HEPES (Invitrogen), 1 μg/ml cyclosporine (CsA) (Sigma), and 10U/mL interleukin-2 (IL-2) (32).

### Neutralization assay

We performed all neutralization assays in duplicate at least three independent times. ART-naïve plasma samples were heat-inactivated at 56°C for 1 hour, and total IgG was isolated from post-ART plasma samples using the Melon gel IgG spin purification kit (Thermo Scientific) according to the manufacturer’s protocol. A 1:50 plasma dilution or equivalent quantity of isolated antibodies was used in the TZM-bl assay. A simian immunodeficiency virus (SIV) Env deleted pseudotyped with vesicular stomatitis virus (VSV) G protein Env was used as a negative control in the neutralization assays. Briefly, heat-inactivated plasma or isolated IgG was incubated with around 1000 infectious particles of a viral strain at 37°C for 1 hour. Approximately 1X10^6^ TZM-bl cells were added to each well with a plasma-virus mix. We estimated the number of infectious virus after two days by quantifying the luminescence in each well using Bright Glo Luciferase Assay System (Promega). Differences in relative luciferase units (RLUs) in the presence or absence of plasma or antibody were used to calculate percent neutralization after subtracting background RLU (TZM-bl cells and growth medium only).

### Antibody-dependent cellular cytotoxicity assay

All ADCC assays were performed in duplicate at a minimum of three independent times. ADCC capacity was assessed against MT4-CCR5-Luc cells infected with different replication-competent variants. Briefly, MT4-CCR5-Luc cells were infected with different virus stocks for four to seven days, and the infected cells were used as targets when RLUs were at least 10-fold over that observed in the MT4-CC5-Luc cells not exposed to a virus. Heat-inactivated 1:50 dilution plasma samples or an equivalent amount of isolated antibodies were added to infected 1X10^5^ MT4-CCR5-Luc cells at 37°C. After 15 minutes, 5X10^4^ of NK (CD16^+^KHYG-1) cells were added at 1:1 dilution to each well (32). After 24 hours, wells were assessed for luminescence using Bright Glo Luciferase Assay System (Promega) per manufacturer’s instructions. Differences in relative luciferase units (RLUs) in the presence or absence of plasma or antibody and growth medium were used to calculate percent ADCC after subtracting background RLU (uninfected MT4-CCR5-Luc cells and CD16^+^KHYG-1 cell in RPMI).

### Cytokine levels

All assays were performed in duplicate. Plasma levels of fifteen cytokines including IL-6, CXCL10, IL-17A, IL-17F, IL-4, IL-2, IL-13, IP-10, CCL5, RANTES, TNF-β, TNF-α, IL-10, IL-12p70, APRIL, BAFF, and sCD40L were measured using the human B cell premix kit from Biolegend Legendplex™ using manufacturer instructions. All analytes were measured on samples that were not previously thawed.

### Immunoglobulins, anti-HSV, and anti-tetanus-toxoid levels

All assays were performed in duplicate. We measured absolute concentrations of IgG, IgG1, IgG2, IgG3, IgG4, IgA, and IgM using a Luminex magnetic-based assay according to the manufacturer’s instructions (Bio-Rad). Plasma levels of Herpes simplex virus 1/2 (HSV-1/2) and *Clostridium tetani* (tetanus)-toxoid-specific IgG were captured and measured with ELISA kits from Calbiotech and Alpha diagnostics according to manufacturers’ protocols.

### Auto-antibody levels

All assays were performed in duplicate. Plasma levels of antibodies reacting to seventeen self-antigens including β-2-Glycoprotein, C1q, CENP-B, Jo-1, Ku, Mi-2, Myeloperoxidase, Proteinase 3, PCNA, PL-12, PM/Scl-100, RNP/SM, Scl-70, Sm, SSA/Ro52, SSA/Ro60, and SSB/La were determined using a magnetic bead panel (Millipore) and quantified on Magpix (Luminex) instrument containing xPONENT 4.2 software (Boston University Analytical Core Facility). Anti-cardiolipin antibody levels were measured by ELISA (Inova Diagnostics) and quantified on BioTek Synergy HT plate reader at 450nm absorbance.

### Single genome amplification (SGA)

Intracellular RNA from participants’ blood samples stabilized in PAXgene BRTs was isolated and purified with PAXgene Blood RNA IVD kit (Qiagen) according to the manufacturer’s instructions. Isolated RNA was reverse transcribed with OFM19 (5’-GCACTCAAGGCAAGCTTTATTGAGGCTTA-3) primer to make viral cDNA. The cDNA was diluted to ensure a maximum of 30 out of 96 PCRs yielded positive reactions using primer sets as protocol as described previously (33, 34). Amplicons from positive PCR reactions were cleaned using ExoSAP IT (Affymetrix) as previously described (31, 35). Viral Envs were sequenced by Sanger-based sequencing (Genewiz) and edited on the Sequencher program (Gene Codes).

### Envelope sequence analyses

HIV-1 Env sequences from all pre-ART participants were aligned using Los Alamos HIV sequence database HIV align tool (https://www.hiv.lanl.gov/content/sequence/VIRALIGN/viralign.html). Sequence alignments were done using the HMM alignment model. A maximum phylogenetic tree was generated using the general-time-reversible (GTR) distance model on the PhyML interface (https://www.hiv.lanl.gov/content/sequence/PHYML/interface.html). The color and symbol-coded phylogram was generated using the rainbow tree tool in LANL (https://www.hiv.lanl.gov/content/sequence/RAINBOWTREE/rainbowtree.html). The mean pairwise Env genetic diversity was estimated based on a GTR substitution model with optimized equilibrium frequencies on DIVEIN as previously described (36). All sequences have been deposited in GenBank (OK513840 - OK514180).

Differential amino acid frequency among PWH/Active Mtb and PWH/No Mtb sequences were determined using the LANL GenSig tool (https://www.hiv.lanl.gov/content/sequence/GENETICSIGNATURES/gs.html). This is a phylogenetically corrected analysis to minimize false positive due to lineage effects. Briefly, the algorithm presents the odds ratio for the differential amino acid prevalence in PWH/Active Mtb as compared to PWH/No Mtb sequences relative to the ancestral state in a phylogenetic tree. We used a false discovery rate (q < 0.1) to correct for multiple comparisons. Among the identified sites, we only considered positions that have previously been associated with changes to broadly neutralizing antibodies (bnAb) susceptibility.

The bnAb-ReP algorithm was obtained from https://github.com/RedaRawi/bNAb-ReP, and the packages were run on the Boston University Shared Computing Cluster (SCC) (37). Input included the predicted amino acid sequence for the 341 Envs in fasta format. The output was probabilities from 0 to 1 with higher values indicating neutralization sensitivity against the 33 different bnAbs.

### Heatmap

The heat maps were generated to highlight neutralization fingerprints in the study using the Los Alamos HIV sequence database heat map tool (https://www.hiv.lanl.gov/content/sequence/HEATMAP/heatmap.html). Euclidean distance method was used to create hierarchical clustering and bootstrap to calculate cluster stability on the heatmap.

### BP score and bnAb-Rep estimation

Breadth and potency (BP) scores for percent neutralization and percent ADCC were estimated as previously described (38). Briefly, BP score consists of an average of the log_2_ transformed % neutralization + 1 or log_2_ transformed % ADCC + 1.

For the bnAb-Rep analysis, each Env – bnAb pair was classified as sensitive or resistant based on the predicted probability above versus below 0.5. A generalized estimating equation (GEE) population-averaged model was used to evaluate differences in chance of sensitivity versus resistance in the PWH/Active Mtb versus PWH/No Mtb Envs for each of the selected bnAbs. The GEE model used a binomial distributed dependent variable (sensitive = 1 and resistant = 0), and the data was grouped according to subject to account for the multiple Envs from single individuals. Within each subject, the Envs were deemed to have an independent correlation structure.

### Statistics

Statistical analyses were performed using Stata (version 17.0) and GraphPad Prism 9.4. Summary data for each cohort are depicted as boxes and whiskers with the median and interquartile range indicated. Unpaired t-tests with Welch’s correction and Kruskal Wallis tests were used to assess comparisons between the groups prior to and after ART respectively. Mann-Whitney test was used to compare groups with the non-parametric distribution. Fisher’s exact test was used to assess proportion differences among groups. A nonparametric Spearman or parametric Pearson correlation test was used to determine associations. In the different multi-variate linear regression models, only prior-deemed important disease and demographic factors were included to limit the number of independent variables. Pre-ART multi-variable model independent variables included Mtb status (categorical variable), gender, log_10_ plasma virus level, absolute CD4 count, and age at enrollment. Post-ART multi-variable models included active Mtb disease versus no disease (either prior Mtb or no Mtb), gender, plasma virus level post-ART, and absolute CD4 count post-ART start as independent variables. The geographic origin of the samples was not included in the multi-variable analysis because of the large number of countries. In all evaluations, we considered two-sided p-values of less than or equal to 0.05 as statistically significant. The Bonferroni correction was used to account for multiple comparisons.

### Study approval

The institutional review boards at Boston University, Infectious Disease Institute, Johns Hopkins University, and Joint Clinical Research Centre in Kampala Uganda approved the studies. The ACTG 5274 samples were obtained after new works concept sheet approval. All participants gave written, informed consent for sample use.

## Acknowledgments

We thank the study participants and the participating study sites. The content is solely the responsibility of the authors and does not necessarily represent the official views of the National Institutes of Health.

## Author contributions

MS conceived and designed the study. BA, YM, AJO, EN, and MZ conducted the experiments. LN, YCM, AG, MCH, JK, and KRJ provided clinical samples, data, and critical input. BA and MS analyzed the data and conducted statistical analyses. BA and MS wrote the manuscript with editorial assistance from the co-authors. The AIDS Clinical Trials Group A5274 (REMEMBER) Study Team provided samples, and Supplementary Table 9 lists the investigators.

